# The role of integrin α_v_ and CD44 in GBM migration in human brain

**DOI:** 10.1101/841726

**Authors:** Zev A. Binder, Sarah Hyun Ji Kim, Pei-Hsun Wu, Anjil Giri, Gary L. Gallia, Carlos A. Pardo, Denis Wirtz

## Abstract

Glioblastoma (GBM) is the most common primary adult malignant brain tumor. Recurrence is driven invading tumor cells that escape surgical resection and demonstrate resistance to standard-of-care chemotherapy and radiotherapy. A large body of research has been conducted on tumor cell motility. However, typical *in vitro* models make use of polystyrene culture dishes, which exhibit significantly different physical parameters than brain tissue. Here we report on the use of human organotypic brain slices as an *ex vivo* approach for the dynamic study of GBM cell motility.

Temporal lobectomy tissue from epilepsy patients was obtained and cut into 350µm thick slices. After the tissue slices had a week’s incubation for recovery, fluorescently labeled tumor cells were seeded. We then tracked individual tumor cells using time-lapse fluorescent confocal microscopy. Quantification of motility characteristics, including mean squared displacement, total path length, and consistency, allowed for comparison of different conditions, including knockdown of cell surface proteins integrin α_v_ (ITGAV) and CD44.

Human organotypics demonstrated minimal variability across specimen in terms of motility parameters, including total path length, averaged instantaneous velocity, and consistency. Knockdown of the traditional motility protein ITGAV showed little effect on overall motility while knockdown of CD44 resulted in a significant reduction in both averaged instantaneous velocity and total path length. When the same parameters were examined using Matrigel, ITGAV and CD44 both showed decreased motility, highlighting the impact of the physical environment on cell behavior. Finally, cell motility in mouse organotypic slices was decreased when compared to human organotypic slices.

Here we demonstrate the use of human organotypic brain slices in the study of GBM cell invasion. This model system offers a physiologically-relevant environment in which to examine the dynamic process of cell motility.

## Introduction

Glioblastoma (GBM) is the most common malignant, primary, adult brain tumor [1]. The current survival of GBMs is 14.6 months with the full scope of treatment, encompassing surgical resection, radiotherapy, and chemotherapy [2]. By their nature, GBMs are invasive tumors. Recurrent tumor after surgical debulking and adjuvant therapy occurs most frequently within the radiation field, suggesting that local invasion of malignant cells away from the tumor bulk are the cause of regrowth and subsequent death [3-5]. This is supported by evidence that most of the causes of death in GBM are related to tumor growth in the brain and not metastasis or systemic causes [6]. Migration and direct invasion of GBM cells is therefore an area of great focus in research.

The majority component in the brain is hyaluronic acid [7]. Elsewhere in the body, the major component of the extracellular matrix is collagen [8]. In the brain, collagen is relatively scarce and mostly associated with blood vessels [9]. Similarly, while ECM components such as vitronectin and fibronectin are found in numerous other locations, in the brain they are confined to blood vessels. This unique ECM makeup plays an important role in tumor behavior in the brain. Appropriate model systems need to take into account these differences.

Current model systems used for GBM research include traditional *in vitro* cell line-based assays and *in vivo* animal studies. *In vitro* model systems offer the advantages of being easy to use, relatively inexpensive, and fast growing. However, these models lack key elements of the pathology they are attempting to model, including the biochemical and biophysical microenvironment and three-dimensional structure inherent to human brain tissue. *In vivo* model systems address these limitations, but have restrictions of their own [10]. Species differences may result in non-applicable results and animal experiments are often not designed like clinical trials. Evidence of the limitations of current GBM models is found in the disparity between basic research findings and successful new treatments for GBMs in the clinic. Despite a large number of basic science publications in the GBM field, the last 15 years has seen only two new treatments approved for GBMs [11, 12].

Here we present an alternative model system for the study of human GBM cell motility and invasion, which features advantages of both *in vitro* and *in vivo* model systems. Using human organotypic brain slices as scaffolding for tumor growth, we explored the dynamic process of GBM cell invasion within human brain tissue. To demonstrate the utility of the model system, we investigated the effects of depletion of integrin α_V_ and CD44 on GBM cell motility. These two cell-surface proteins have been identified to have key functions in GBM cell motility, but only the loss of CD44 demonstrated an effect on tumor cells in organotypic slices. Finally, we compared motility results from cells in the brain slices to those from cells growing on standard Matrigel and in mouse brain organotypics, highlighting the importance of species-specific tissue.

## Materials and Methods

### Cell lines

The patient-derived GBM oncosphere cell line JHH-136 was established as previously described [13]. Briefly, tissue was obtained from the Johns Hopkins Hospital (JHH) operating rooms under IRB approval and, within 1-h resection, was dissociated mechanically and enzymatically. Following a red blood cell lysis and centrifugation through a sucrose gradient, the resulting cells were cultured in NeuroCult medium (StemCell Technology, Vancouver, BC, Canada) supplemented with 0.2% heparin (StemCell Technology), hEGF (20 ng/mL, Peprotech, Rocky Hill, NJ), hFGF-b (10 ng/mL, Peprotech), and 1% penicillin/streptomycin. Cell line uniqueness was verified using short tandem repeat profiling through the Johns Hopkins Genetic Resources Core Facility.

### Brain slices

Brain tissue was obtained from temporal lobectomy cases at JHH under IRB approval. Approximately 1cm^3^ sections were sliced into 350-µm thick axial slices on a McIlwain Tissue Chopper (Mickle Laboratory Engineering Co., UK) incorporating both gray and white matter. Sections were placed on MilliCell Inserts (EMD Millipore, Billerica, MA) over 300 µL of organotypic medium. Organotypic medium was comprised of 50% MEM (Sigma-Aldrich, St. Louis, MO) plus 1.2% HEPES (Sigma-Aldrich), pH 7.3, 25% HBSS (Gibco, Grand Island, NY) plus 2.6% glucose (Sigma-Aldrich), 25% horse serum (Atlanta Biologicals, Flowery Branch, GA), and 1% l-glutamine (Lonza, Allendale, NJ). Medium was changed every two to three days. No additional actions were taken for one week in order to allow the inflammatory response caused by slicing to subside.

### Lentiviral transduction and virus production

shRNA sequences for ITGAV (integrin α_v_) and CD44 were obtained as glycerol stock in pTRIPZ inducible plasmids (Thermo Scientific). For transduction, DNA plasmids were transfected into Oneshot Stbl3 Competent Cells (Invitrogen) using the heat shock method. After a short incubation period in ice, a mixture of competent bacteria and DNA was placed at 42°C for 45 s and then placed back in ice. Transformed bacteria were then incubated in SOC medium at 37°C in a shaker for 1h. Bacteria were next plated on an agar plate for 16 h. After incubation, one colony of the bacteria was transferred into a larger volume of LB medium and inoculated overnight. After the inoculation, DNA plasmids were collected from the competent bacteria using MidiPrep (Clontech, Mountain View, CA).

Viruses containing the plasmids of interest were produced by transfecting 293T cells. A mixture of VSVG, ΔR8.91, and DNA plasmids were combined with 2M CaCl_2_. Next, 2X HBS solution was added into the mix, which was then added to the cell plates. Medium was collected on the second and third days of the transfection, then ultracentrifuged at 20,000g at 4°C for 3 h to concentrate the virus. Viral pellets were then resuspended in 500 µL of NeuroCult medium.

### Cell transfection

GBM oncospheres were passaged to dissociate the spheres into single cells. Three to five days after passaging, concentrated virus was added to the cells along with 8 ng/mL polybrene (Sigma-Aldrich) and the cells were incubated at 37°C plus 5% CO_2_ in humidified air. 24 h after transfection, medium was changed. In order to induce tRFP expression, Doxycyline (Santa Cruz Biotechnology, Dallas, TX) was added to the culture for a final concentration of 4 µg/mL daily for three days. The cells were then subjected to cell sorting to isolate an RFP-positive population. The analysis and sorting were performed using Summit 4.3 software (DAKO USA, Carpinteria, CA) on a MoFlo MLS (Beckman Coulter, Brea, CA) sorter equipped with a Coherent Enterprise II 621 laser.

### Flow cytometry

Cells were triturated, counted, and 100,000 cells aliquoted per vial. After incubation with human serum (Sigma-Aldrich) for 15 min, the cells were washed in PBS + 1% BSA. The cells were then incubated with APC-tagged primary antibody, either anti-α_V_β_3_ (R&D Systems, Minneapolis, MN), anti-CD44 (BD Pharmingen, San Jose, CA), or APC-tagged isotype antibody, IgG_1_ for α_V_β_3_ (R&D Systems) and IgG_2B_ for the CD44 (BD Pharmingen) for 20 min. They were washed twice with PBS + 1% BSA before incubating with 7-Amino-actinomycin D (7AAD) (BD Biosciences) for 10 min. Cells were run on a FACSCalibur (BD Biosciences) and data collected using CellQuest. Live cell determination was conducted using 7AAD positivity. Analysis was conducted using FlowJo (Ashland, OR) to determine degree of knockdown. Mean fluorescence was corrected using the isotype staining and knockdown was calculated by the decrease in mean fluorescence between the control cells and the knockdown cells.

### Embedding MEF cells in 2D collagen I matrix and cell migration

Plates were coated with 50 µg/mL of type I collagen (BD Biosciences) using 12-well plates. Plates were incubated at 37°C for 1 h and then washed 3 times with PBS. Mouse embryonic fibroblast (MEF) cells were added at a concentration of 2,000 cells/mL and given 24 h to attach and grow prior to imaging.

### Embedding HT1080 cells in 3D collagen I matrix and cell migration

Gel formation and cell embedding was performed as described previously[14]. Briefly, 2 mg/mL type I collagen gels were used with HT1080 fibrosarcoma cells at a concentration of 36,000 cells/mL, to be able to track single cells. Gels were incubated for 24 h at 37°C supplemented with 5% CO2 prior to imaging.

### Cell motility in Matrigel

Matrigel (BD Biosciences) was diluted to 250 µg/mL, added to a glass-bottomed plate, and incubated at 37°C for 1 h. Cells were spun down, triturated, counted, and 10,000 were resuspended in 500 µL of NeuroCult medium. The medium-cell mixture was placed on top of the Matrigel coating and the dish was placed in the incubator. Medium was changed every 2-3 to days. Three days prior to imaging, 4 µg/mL of doxycycline was added to the cells to induce shRNA and tRFP expression.

### *Ex vivo* tumor setup

GBM oncosphere cells were spun down, triturated, counted and 20,000 cells per brain slice were resuspended in 3 µL per brain slice of NeuroCult medium minus penicillin/streptomycin. Medium was changed to a 1:1 mixture of organotypic medium to NeuroCult medium and 3 µL of tumor cells were added to each slice. After 1 to 4 days of tumor growth, 4 µg/mL of doxycycline was added to the medium below each slice each day to induce shRNA and tRFP expression. After three days of doxycycline treatment, the tumor slices were placed on the microscope for imaging.

### Microscopy

For the brain slices, time-lapse fluorescence and differential interference contrast (DIC) imaging were obtained using an A1 confocal microscope (Nikon, Tokyo, Japan) with NIS-Elements Software (Nikon). Images were acquired using a 10x objective with images taken every 10 min for a total of 24 h on two to six positions per brain slice. A Z-stack of 50 µm was obtained at each position and final analysis was conducted using a maximum-intensity projection. For the MEFs and HT1080s, cells were imaged every 2 min for 16.5 h using an ORCA-AG 1K CCD camera (Hamamatsu Photonics) mounted on a TE 2000 microscope base with NIS-Elements software (Nikon).

### Cell tracking and analysis

Organotypic tumor cell tracking was conducted manually using the tracking package in NIS-Elements software. MEF and HT1080 cell tracking was conducted using a template match algorithm in Metamorph imaging software (Molecular Devices, Sunnyvale, CA) From the cell positions, velocity, path length, distance from objective, Mean-squared displacement (MSD), and alphas were calculated using MS Excel and custom software. Cell density in brain slices was obtained using NIS-Elements.

## Results

### Dynamic imaging of human GBM tumor cells in human brain slices

We initially demonstrated the feasibility of our human organotypic brain slice model system using tRFP-labeled GBM tumor cells (Fig. 1). We used lateral cortex tissue from the temporal lobe as this tissue was readily available from surgical resections on patients with medically-intractable epilepsy. To maintain tissue viability, we minimized the time between removal of the tissue and placement in a humidified incubator supplemented with 5% CO_2_. The tissue was obtained in approximately 1 cm^3^ pieces and sliced axially into 350 µm thick slices. This thickness was thin enough to allow for passive diffusion of nutrients, but thick enough to immerse cells in a 3D environment [15]. To avoid confounding effects from the tissue response to the trauma of slicing, brain slices were given a week in the incubator to recover. After recovery, 20,000 GBM oncosphere cells in a single-cell suspension were placed on the top of each slice, in 3 µL of NeuroCult medium without penicillin/streptomycin. The GBM oncosphere cells were a patient-derived cell line established at JHH, cultured in medium selecting for tumor stem cell growth. The tumor cells were given 72 to 96 h to establish themselves in the brain tissue before 4 µg/mL of doxycycline was added to induce expression of the shRNA sequences. Tissue slices were then placed on an inverted confocal microscope; time-lapse fluorescence and DIC images were collected every 10 min for 24 h.

**Figure 1.**
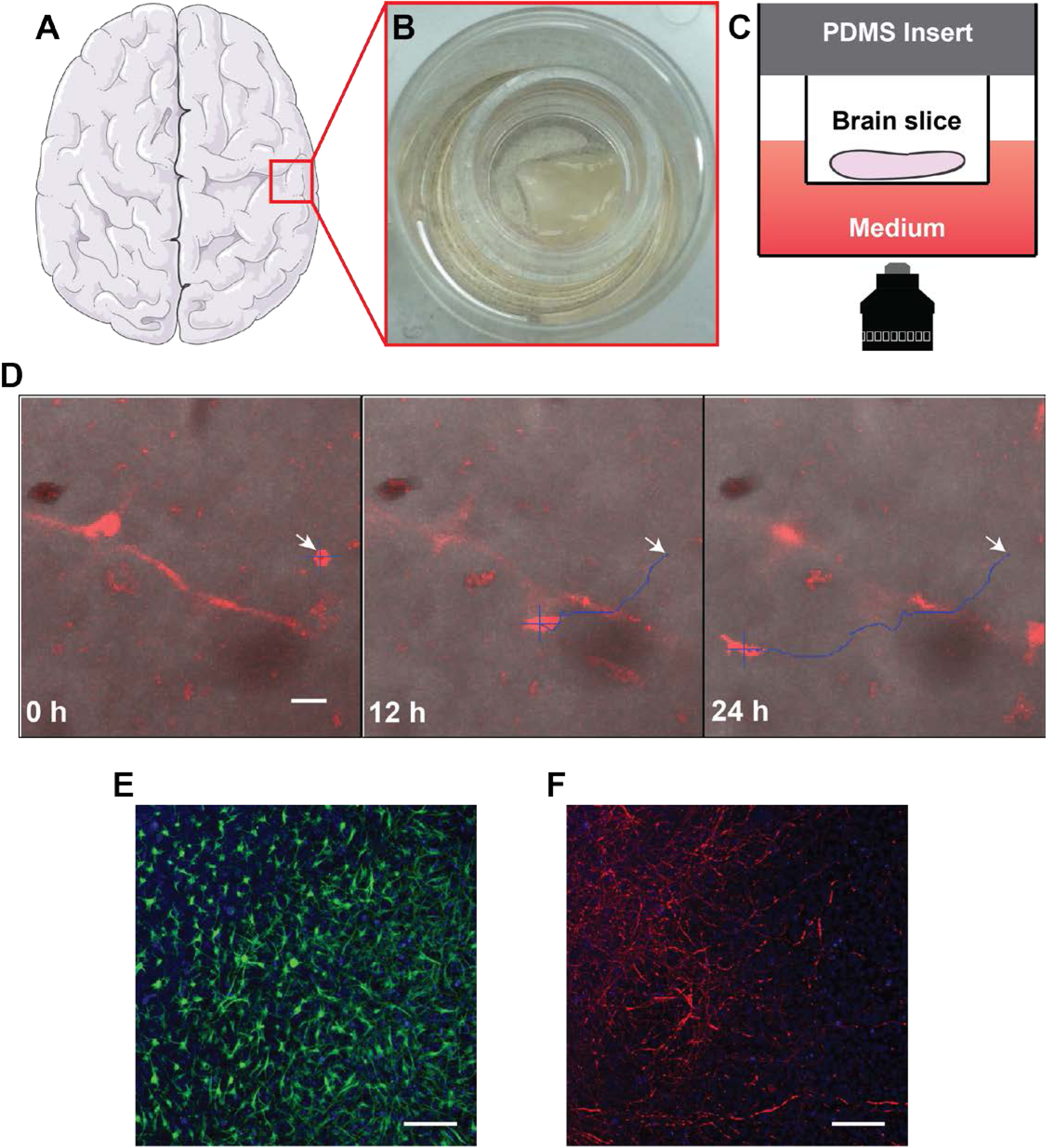
Organotypic brain slice experimental setup. (A) Tissue slices are obtained from temporal lobectomy surgeries and maintained in a humidified environment at 37°C with 5% CO_2_ providing live tissue slices for further experimentation (B). After tumor implantation the slices are placed on an inverted confocal microscope (C) for live cell fluorescent imaging across 24 hours (D). The white arrow marks the cell’s origin, the blue plus marks the cell body, and the blue trajectory shows the cell’s path. (E-F) Immunofluorescence image of brain slices showing GFAP staining for astrocytes (E) and Tuj1 staining for neurons (F). Scale bar, 50 µm.

A variety of motility parameters describing the ability of GBM cells to migrate and invade the brain were computed from the time-dependent positions of the cells. From the coordinates of cell centroids, population-averaged instantaneous velocity, total path length, and consistency, as defined by final distance from the origin divided by total path length, were calculated (Fig. 2). The mean instantaneous velocity was 0.09 µm/min, with a wide spread between 0.02 µm/min and 0.35 µm/min. For comparison, fibroblasts crawling on a flat collagen substrate and fibrosarcoma cells moving within a 3D collagen matrix move with a mean speed of 0.68 µm/min and 0.33 µm/min, respectively (Supplementary Fig. 1). Interestingly, we found that variation in cell speed for GBMs in human slices was significantly larger than for the other two model systems as measured by the coefficient of variation (CV). The CV of cell speed for GBMs in human brain slices in our measurement was 1.4 in comparison to value of 0.9 from both the other two models. We next measured how consistent the moving orientation is for the GBM cells in human brain slice. The Consistency was measured by dividing final distance from origin by total path length, resulting in a value between 0 (no consistency, cell move at random orientation) and 1 (highly consistency i.e. cell keeps moving at the same direction at all time) and the GBM cells shows a mean consistency value of 0.23 with range spreading from 0.02 to 0.72.. This indicate GBM cells showed little consistency in their migratory patterns, as indicated by low consistency values (Fig. 2).

**Figure 2.**
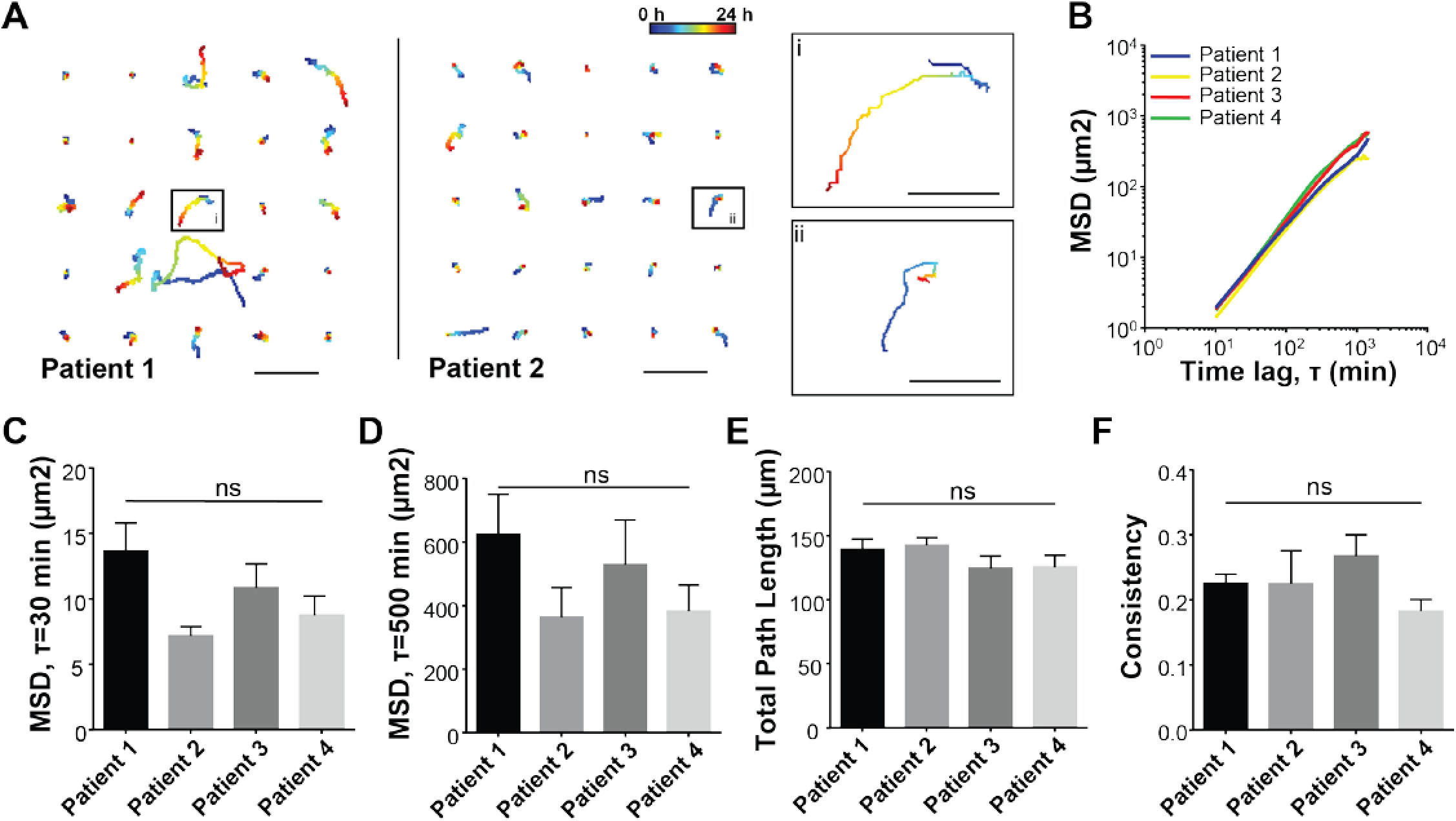
Reproducibility of the organotypic model system. (A) Trajectories of 25 randomly selected cells from Patients 1 and 2, highlighting cell heterogeneity and lack of global symmetry. Scale bars, 100 µm; inset scale bars, 25 µm. Movies of migrating cells were recorded for 24 h at a frame rate of 1 frame per 10 min. Cells were seeded on patient brain sections at an initial density of 20,000 cells per slice. (B) Population-averaged MSDs of cells for the four patients showing significant overlap. (C-D) Mean squared displacement (MSD) values at 30 min (C) and 500 min (D) showing no significant difference between the four patients. (E-F) Total path length (E) and consistency (F) as measured by final distance from the origin divided by the total path length, demonstrating no significant difference between the four patients. ns: non-significant.

Its known that the cell migration under no external driving force (such as chemo-attractants) commonly exhibit persistent random walk (PRW) patterns [16]. To examine whether the GBM cells migration in human brain slices was described by PRWs we calculated the MSD from the trajectories of the cells. The MSD profile from GBM model featured a slope greater than one at short time lags and approaching one at long time lags. These results suggest that the motility patterns of GBMs cells in human brain slices display characteristics of the PRW model and that, at short time lag, the cell movement is persistent (MSD slope >1) while at long time lag the cell movement is random (MSD slope =1) [16]. Of note, the trajectories did not display an isotropic distribution as would be expected with cells following a persistent random walk (Supplementary Fig. 2).

### Reproducibility of the organotypic model system

At least three brain slices, each from four different patient samples, were characterized to assess the reproducibility of our model system (Fig. 2B [better be specific to subpanel level]). The patients’ ages ranged from 5 to 29 years old (Table 1); all four patients were female. Each patient had a different underlying cause leading to the temporal lobectomy surgery, although all had epilepsy. The tissue, although always taken from the temporal lobe, was taken from different locations within the temporal lobe. Despite these differences in organotypic tissue, we found no statistically significant differences in motility parameters, including population-averaged instantaneous velocity, total path length, consistency, and MSDs evaluated at different time lags for these brain slices. MSD curves overlapped with non-significant differences at all probed time lags. This result indicates that despite wide cell-to-cell variability in migration (Fig. 2A), we observed high consistency of GBM migration across different patient brain slices (Fig 2F).

**Table 1.** Demographic and timing data of organotypic brain slices.

|  | **Patient 1** | **Patient 4** | **Patient 2** | **Patient 4** |
| --- | --- | --- | --- | --- |
| **Age (Years)** | 29 | 5 | 13 | 13 |
| **Gender** | F | F | F | F |
| **Tissue Age (Days)** | 21 | 12 | 7 | 11 |
| **Tumor Age (Days)** | 6 | 5 | 3 | 7 |

Next, we assessed whether the local cell density in the brain or the density of GBM cells influenced the speed of individual GBM cells. Densities of both normal cells and tumor cells were calculated at the beginning of each video in a diameter of 2.5x and 5x the average cell diameter (=16.3 µm) around each tracked cell. The density of tumor cells and non-tumor cells did not predict the instantaneous velocity (Supplementary Fig. 3). Together, these results demonstrated the robustness of our model system.

### Depletion of integrin α_V_ induces no significant change in motility

To demonstrate the usefulness of our organotypic model system, we next investigated the effect of depletion of two key cell surface proteins on GBM motility. ITGAV was chosen as a specific ITGAV inhibitor, Cilengitide, previously underwent Phase II and III clinical trials for GBMs. ITGAV depletion through shRNA resulted in an average loss of 37% of ITGAV expression, determined by analytical flow cytometry (Fig. 3A). Cells behaved similarly regardless of expression of ITGAV, moving with similar speed and over a similar distance. Differences in total path length were non-significant (Fig. 3F).

**Figure 3.**
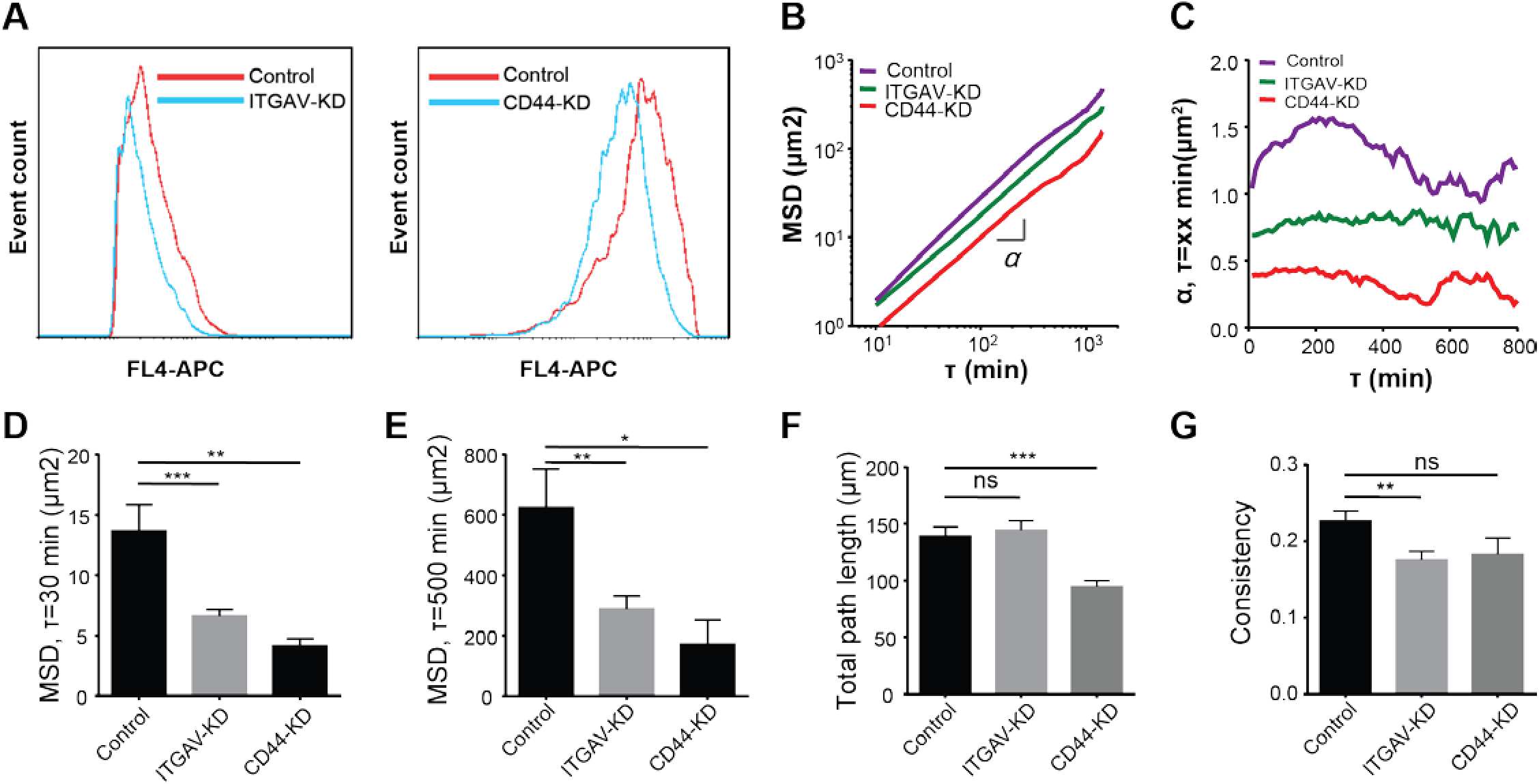
Depletion of ITGAV induces no effect on cell migration, while depletion of CD44 decreases cell migration. (A) Flow cytometry histogram demonstrating the depletion of ITGAV (left) and CD44 (right). (B) Average mean squared displacement (MSD) for control, ITGAV-depleted, and CD44-depleted cells. (C) Alpha values obtained from fits of MSDs with a power law of exponent alpha for control, ITGAV-depleted, and CD44-depleted cells. (D-E) MSD values measured at (D) 30 min and (E) 500 min time lags. (F) Total path length and (G) consistency as measured by final distance from origin divided by total path length. Each slice was imaged at four to six different positions for 24 h, with at least two different patients’ tissue used per condition. ns: non-significant; *: p ≤ 0.05; **: p ≤ 0.01; ***: p ≤ 0.001; ****: p ≤ 0.0001.

Tracking of ITGAV-depleted cells again demonstrated no degree of coordinated movement (Fig. 4). Consistency was significantly decreased suggesting a lesser degree of directed movement. The mean value for the ITGAV-depleted cells was 0.18 compared to 0.23 for the control cells. In support of this data, exponent α for MSDs were close to unity regardless of time lag, which indicates movement akin to pure diffusion. These elevated α suggest directed movement, while the smaller α for the ITGAV-depleted cells point towards restricted movement. Cell heterogeneity was not significantly affected by ITGAV depletion, except in the case of instantaneous velocity (Supplementary Fig. 4). Taken together, these results indicated that the depletion of ITGAV leads to a small effect on GBM cell motility.

**Figure 4.**
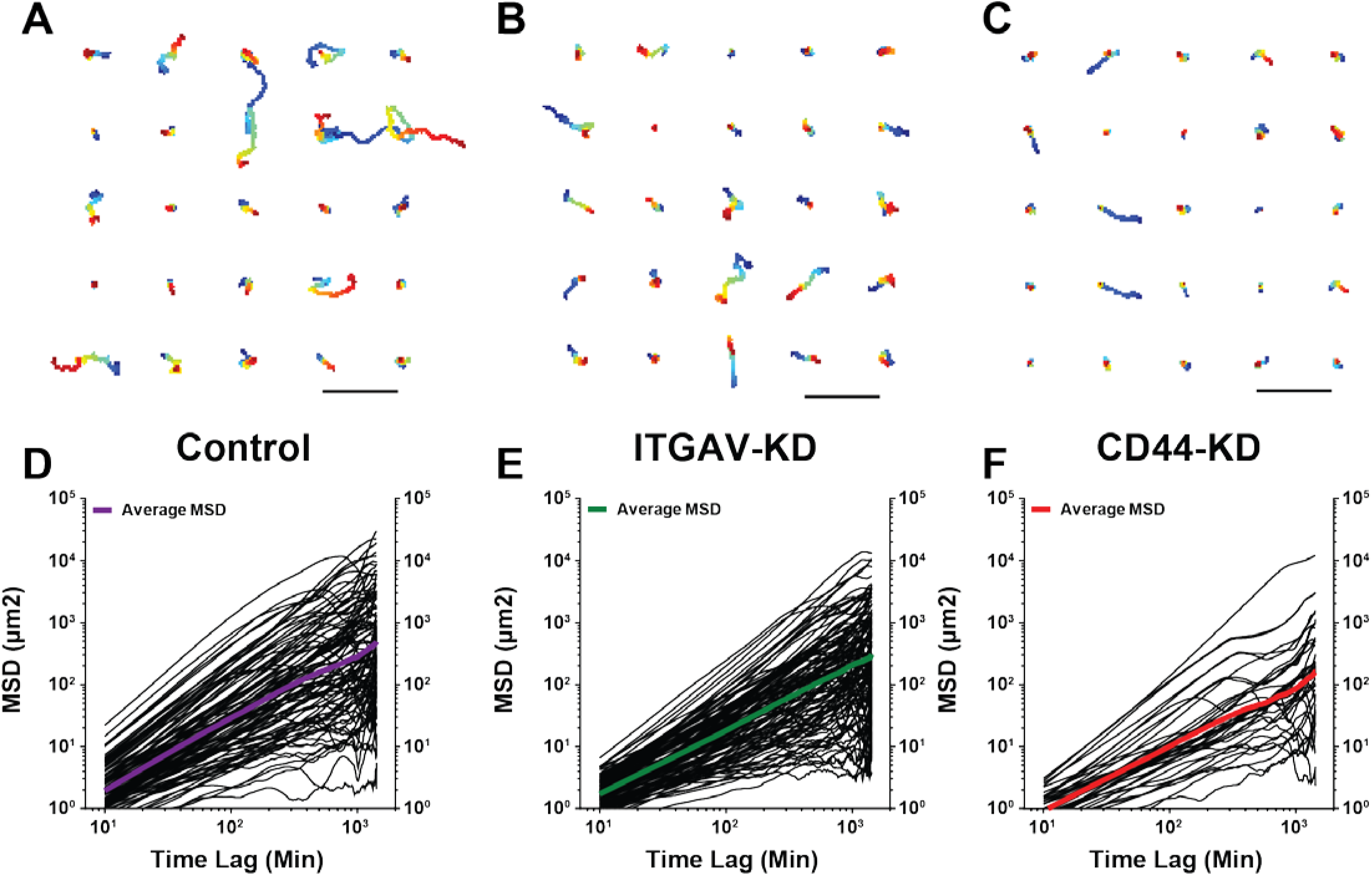
Tumor cells exhibit a large degree of heterogeneity. (A-C) Trajectories for 25 randomly selected cells from (A) control, (B) ITGAV-depleted, and (C) CD44-depleted cells. Scale bars, 100 µm; inset scale bars, 25 µm. (D-F) MSDs of (D) control, (E) ITGAV-depleted, and (F) CD44-depleted cells.

### Depletion of CD44 induces reduction in motility

As the main ECM targets of ITGAV, vitronectin and fibronectin, are not found in high concentrations in the brain, we chose CD44 as our second target protein. CD44 binds to hyaluronic acid, the main component of brain ECM. shRNA-induced depletion of CD44 induced significant decreases in multiple aspects of cell motility (Fig. 3A). Population-averaged instantaneous velocity was decreased by 21% in the CD44-depleted cells compared with control cells. Similarly, total path length was decreased by 19% in the CD44-depleted cells compared to control cells. Consistency was not statistically significantly different with a mean value of 0.18 in the CD44-depleted cells, compared to 0.23 for the control cells. MSDs were significantly decreased at all time lags (Fig. 4). For instance, the mean MSD value at 30 min time lag for the CD44-depleted cells was >70% shorter than for control cells. Cell population heterogeneity was not affected for any of the quantified motility parameters (Supplementary Fig. 3).

These results highlighted the significant effect of CD44 depletion on cell motility in human brain. Population-averaged instantaneous velocity and total distance traveled were both decreased while consistency seemed not to be affected.

### Comparison of brain slice motility to Matrigel motility

GBM oncosphere cells moved significantly faster and farther on Matrigel coating than in brain tissue slices (Fig. 5). Trajectories visually demonstrated the longer paths cells took on Matrigel when compared to human brain tissue. These findings were confirmed upon quantification of motility parameters. Control cells on Matrigel-coated plates moved at a faster mean instantaneous velocity, nearly twice that of cells in brain slices. The mean path length on Matrigel was more than twice as long as in brain slices.

**Figure 5.**
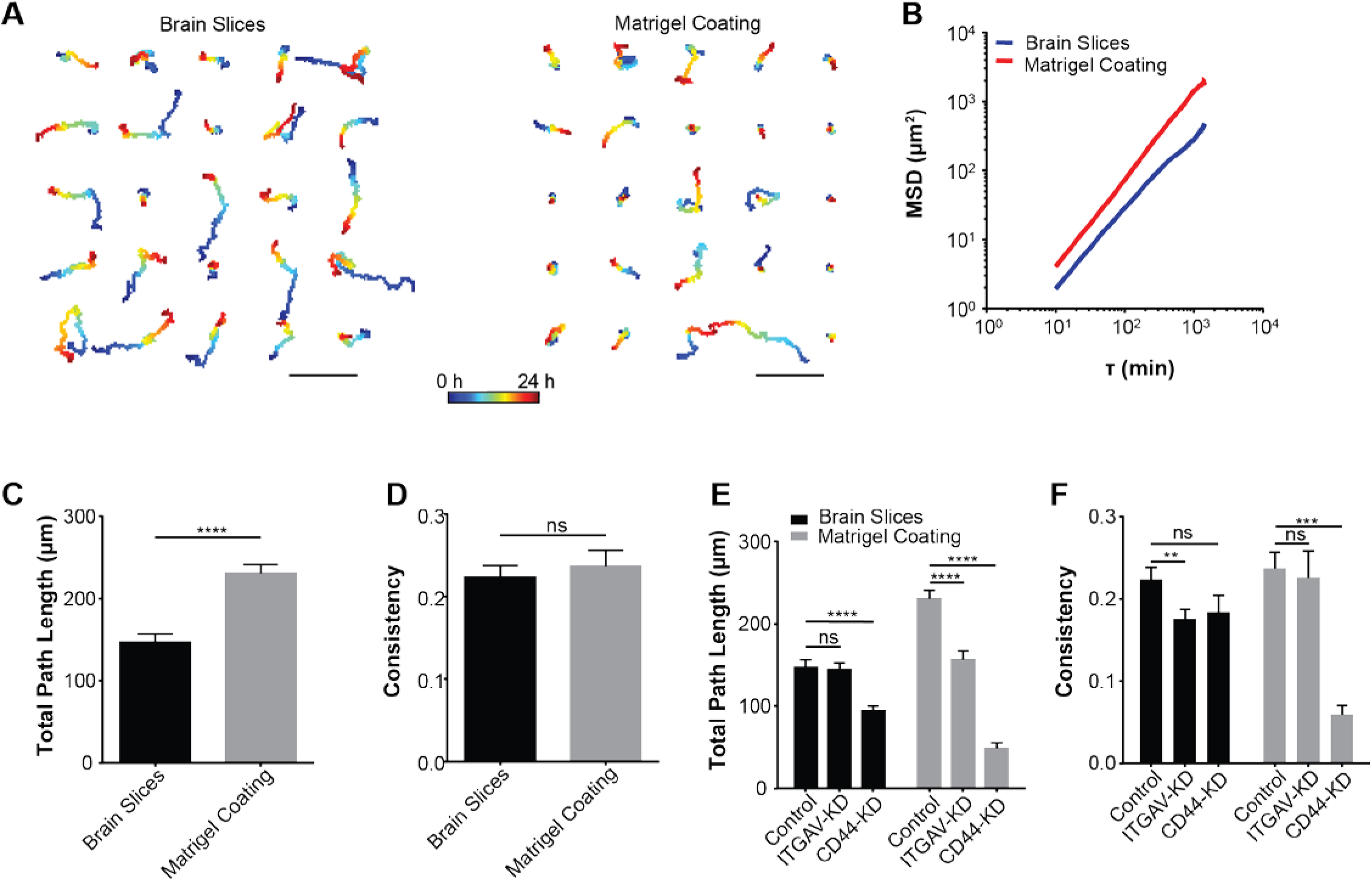
Comparison of motility descriptors on Matrigel *vs.* in brain slices. (A) Trajectories of 25 randomly selected cells from brain slices (left) and Matrigel coating (right). Scale bars, 100 µm; inset scale bars, 25 µm. (B) Population-averaged mean squared displacement values for brain slices (blue) and Matrigel coating (red). (C-D) Motility parameters showing the difference in cell migration in brain slices *vs.* on Matrigel coating. (C) Total path length and (D) consistency as measured by final distance from origin divided by total path length. (E-F) Contrasting the effect of ITGAV depletion on motility characteristics in brain slices compared to on Matrigel coating. (E) Total path length and (F) consistency as measured by final distance from origin divided by total path length. Initial cell densities for the Matrigel coated plates were 10,000 cells per well. Each well was imaged at 4-6 different positions for 24 h. ns: non-significant; *: p ≤ 0.05; **: p ≤ 0.01; ***: p ≤ 0.001; ****: p ≤ 0.0001.

The effect of ITGAV depletion on GBM oncosphere cells was markedly different in cells moving on Matrigel compared to cells in brain tissue slices. On Matrigel, depletion of ITGAV induced a significant reduction in the instantaneous velocity and total path length compared to the control cells. Consistency was unchanged by depletion of ITGAV. In each case, the effect was the contrary to what was found in the brain tissue slices.

Averaged instantaneous velocity and total path length were significantly decreased with depletion of CD44, and to a much larger degree than in the brain slices. There was ∼80% decrease in both the instantaneous velocities and the total path length. Consistency, while unchanged in the brain slices with CD44 depletion, showed a significant reduction of 75% from the control cells on Matrigel. These results demonstrated the strong effect the environment has on GBM cell migration. Motility parameters displayed striking differences between in organotypic slices and on Matrigel.

### Comparison to mouse brain slices

To compare the motility of GBM oncosphere cells in human brain slices to mouse brain slices, tumors were grown in mouse brain slices and subjected to the same imaging method and motility analysis. Mouse brains demonstrated reproducibility with three different mouse brains providing non-significantly different values for population-averaged instantaneous velocity, total path length, and consistency (Fig. 6).

**Figure 6.**
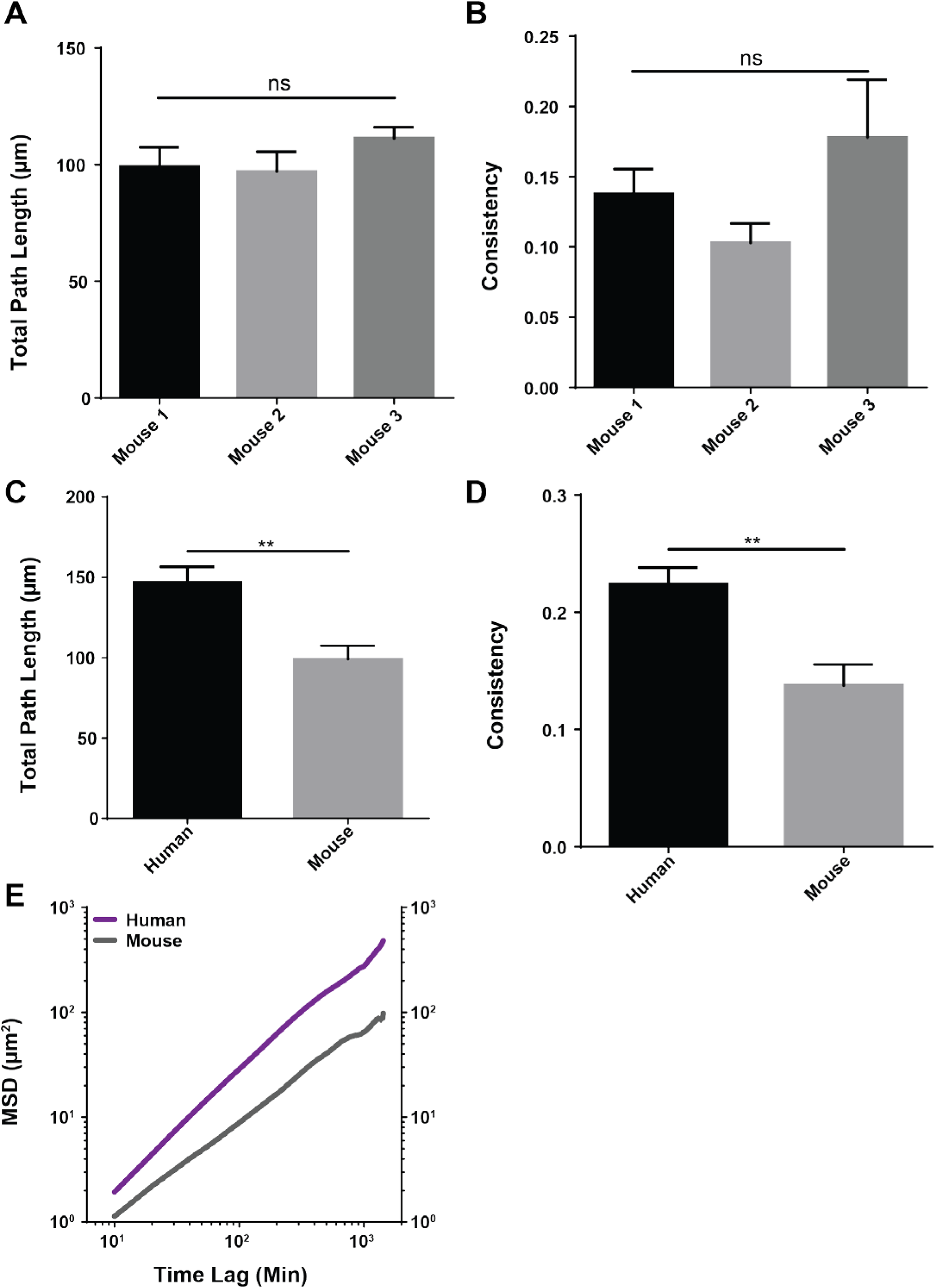
Suspension GBM cell motility in human brains is greater than in mouse brains. (A-B) Motility parameters in mouse brains were found to be reliable and reproducible across multiple samples. (A) Total path length and (B) consistency all demonstrated non-significant changes between 3 mouse brains. (C-E) Comparison of motility parameters in human brain slices to mouse brain slices revealed significant decreases in all parameters. (C) Total path length and (D) consistency all showed between a 25% and 40% decrease in the mouse brains compared to the human brains. (E) Population-averaged mean squared displacement for human and mouse brains.

The mouse brains consistently demonstrated lower values of motility in each category quantified. Population-averaged instantaneous velocity was decreased by > 25% in the mice brains when compared to human brains. Total path length was decreased by nearly a third and the measure of consistency was decreased by just < 40%. Taken together these results highlight that human GBM oncosphere cells do not behave identically in different species’ brain tissue slices.

## Discussion

In this study we demonstrated the viability of time-lapse imaging using GBM oncospheres in *ex vivo* brain slices for the study of GBM cell motility. Model systems that have a higher degree of fidelity to the relevant physiologic microenvironment produce results that are more translatable to the clinic. By using actual human tissues, we circumvented many of the existing issues in traditional *in vitro* culturing, such as dimensionality (2D instead of 3D), microenvironment, other cell types, and extracellular matrix composition.

To show the viability of this model system we tracked the migration of human GBM cells in brain tissue organotypics obtained from temporal lobectomies for epilepsy patients [17]. Tumor cells moved through the existing brain parenchyma, encountering the cell types and structures typically found in the brain. Individual experiments were run out beyond 24 h, demonstrating the flexibility of the system in terms of temporal range.

Hyaluronic acid is an anionic glycosaminoglycan found throughout the body [18]. Its negative charge allows it to attract a large number of cations such as Na^+^ [19]. These in turn exert an osmotic pressure and draw water along with them, giving the brain its spongy plasticity. The Young’s modulus for brain tissue has been experimentally reported at 0.24 to 3.42 kPa [20] and by modeling reported at 9.21 kPa [21]. For comparison, bone, of which collagen makes up close to 90% of the organic matter [22], has an experimentally determined Young’s modulus around 14 GPa, a 10^6^-fold increase in stiffness [23]. Elasticity is known to play a significant role in stem cell differentiation [24], while stiffness modulates GBM cell motility [25]. The physiologic nature of organotypic slices allows for experimentation in the appropriate physical environment.

Brain tissue also has a 3D aspect, which is overlooked in traditional 2D cell culture models (i.e. flat dishes). Work done with cell spheres has shown this 3D environmental impact with decreased availability of target molecules in the center of the spheres leading to decreased effect [26]. In addition to creating a gradient of introduced molecules, 3D structure results in a hypoxic gradient. Hypoxia affects activation of hypoxia inducible factor-1, which in turn leads to the expression of a large number genes affecting vasculature formation and cell proliferation [27].

A third aspect of physiologic brain tissue that traditional cell culture lacks is the presence of the background cellular parenchyma, such as neurons, microglia, macroglia, and vasculature. These normal cells influence tumor growth and motility as it is known that GBM tumor cells have a preference for following vasculature [28] and neuronal axons [19]. Brain slices studies using rat models showed GBM tumor cell movement along cerebrospinal fluid pathways and with an inclination for the white matter [29]. Inclusion of these components in the model system results in behavior that is likely similar to what occurs pathologically in the brain.

*In vivo* models are able to recapitulate many of the above listed characteristics. Limitations with these types of models include cost and temporal limitations. Additionally, there are roadblocks for data acquisition of dynamic processes such as invasion and cell motility. Live animal imaging faces several logistical restrictions to maintain the viability of the animals. Of particular importance for cell motility is the amount of time an animal can be immobilized on a microscope stage for imaging. Due to hypothermia and dehydration risk, these experiments typically do not last longer than 2 h at a time [30].

To demonstrate the robustness of our system and reproducibility with multiple patients’ tissue samples, we performed the same experiments and analysis using brain tissue from four different patients. The resulting motility characteristics showed no significant difference between the four samples. This demonstrated not only the reproducibility of the system but also the robustness of the inherent motility characteristics of the GBM oncosphere cell line. Each tissue slice provided a similar enough environment for the cells to behave in a comparable fashion.

ITGAV was chosen for investigation, as it is the target of the anti-integrin peptide Cilengitide, a drug that has been in a number of clinical trials [31-34]. Cilengitide showed initial promise in *in vitro* studies for treatment of GBMs [35, 36]. Based on those studies, clinical trials in both recurrent and primary GBMs were undertaken [36-39]. The most recent, phase III, clinical trial in primary GBMs, looking at both progression-free survival and overall survival, found no significant changes in these measures [40].

The second protein we investigated was CD44. CD44 is also a cell surface protein but one that binds to hyaluronic acid, the major component of brain ECM [41]. We postulated that loss of CD44 expression would result in a significant reduction in GBM cell motility. This hypothesis was confirmed by the significant reduction in instantaneous average velocity and total path length with the loss of CD44. CD44 inhibitors are not currently in clinical trials for GBMs but there is a Phase II trial for any solid tumor investigating the pharmacokinetics and safety of a monoclonal anti-CD44 antibody, RO5429083 [42]. This suggests that there may be future therapeutic options for CD44 inhibition in gliomas.

Depletion of both ITGAV and CD44 did not significantly reduce the heterogeneity of the motility parameters we investigated, with the exception of population-averaged instantaneous velocity in the ITGAV-depleted population. Often, knockdown and subsequent selection results in a homogenous population introduced artificially. Use of the tRFP expression to select for positively transduced cells circumvented this bottleneck. Given that GBM tumors often exhibit a wide variety of phenotypes within the same tumor [43], retention of this feature heterogeneity is a pathologically-relevant characteristic.

Matrigel, comprised of a number of ECM components including laminin, collagen IV, and entactin, is frequently used in glioma *in vitro* research for motility and invasion assays [44-48]. Comparison of control cells in the two microenvironments revealed a significant difference in the motility parameters with cells on Matrigel moving significantly faster than those in brain slices.

Of particular interest is a comparison of results from human organotypic slices and rodent organotypic slices. Rodent brain, in particular from rats and mice, is more readily available than human tissue, especially considering that not all researchers have access to a large-volume neurosurgery center such as the Johns Hopkins Hospital. Several research laboratories have made use of rat or mouse brain slices and time-lapse imaging to investigate other aspects of GBM cell motility [28, 49-55]. To examine the applicability of using mouse brains in place of human brains, we performed comparison of general motility characteristics in wild type JHH-136 cells grown in human brain slices to mouse brain slices.

Motility quantification revealed that mouse brain slices are as reproducible as human brain slices. However, rigorous comparison revealed statistically significant migratory differences between the two species. In instantaneous velocity, total path length, and consistency, the mouse brains presented overall lower values. Given the ubiquitous role of rodent models in GBM studies, it is important to keep in mind the limitation that interspecies differences will affect the experimental results. A possible explanation for the decreased motility parameters in the mouse brains is the difference in cell densities between the species. Previous work has shown that the synaptic density in rodents is higher than in humans [56]. Higher cell density may lead to decreased porosity and decreased porosity has been shown to decrease cell spreading and migration [57].

Several groups have made use of human brain tissue in organotypic slice experimentation, focused on utilizing tumor tissue instead of normal brain [58, 59]. A recent report presented a method for agarose embedding of tissue slices for cutting by a horizontal, vibratome blade [59]. Overall velocities of GFP-labeled GBM cells showed similarities to our results, suggesting differences in cutting methods, using either a horizontal or vertical blade, may not cause significant differences in tissue architecture. Additional work comparing neurosphere dispersal in mouse, pig, and human brain tissue demonstrated no significant differences between the substrates [58]. Of note, the brains were fixed in paraformaldehyde or formalin prior to *ex vivo* culturing, raising the concern that similar outcomes are a result of the fixation process and not the species’ physiological similarities.

In this study we have demonstrated the viability of using human organotypic brain slices combined with GBM oncosphere cell lines to observe and quantify motility and invasion characteristics. This approach has shown robustness by displaying statistically non-significant results upon repeated assays in multiple patients’ brain tissue specimens. The use of this model system has been validated through the investigation of the roles of ITGAV and CD44 in GBM cell motility. We reported a lack of noteworthy effect on motility with the depletion of ITGAV and a significant reduction in motility with depletion of CD44. Finally, we demonstrated the potential confounding effect due to experimental environment by highlighting the different results seen with ITGAV depletion on Matrigel compared to in brain slices. Taken together, these results showcase the usefulness of the human organotypic brain slice model system for study of GBM cell motility.

## Supplementary figures

**Supplementary Figure 1.**
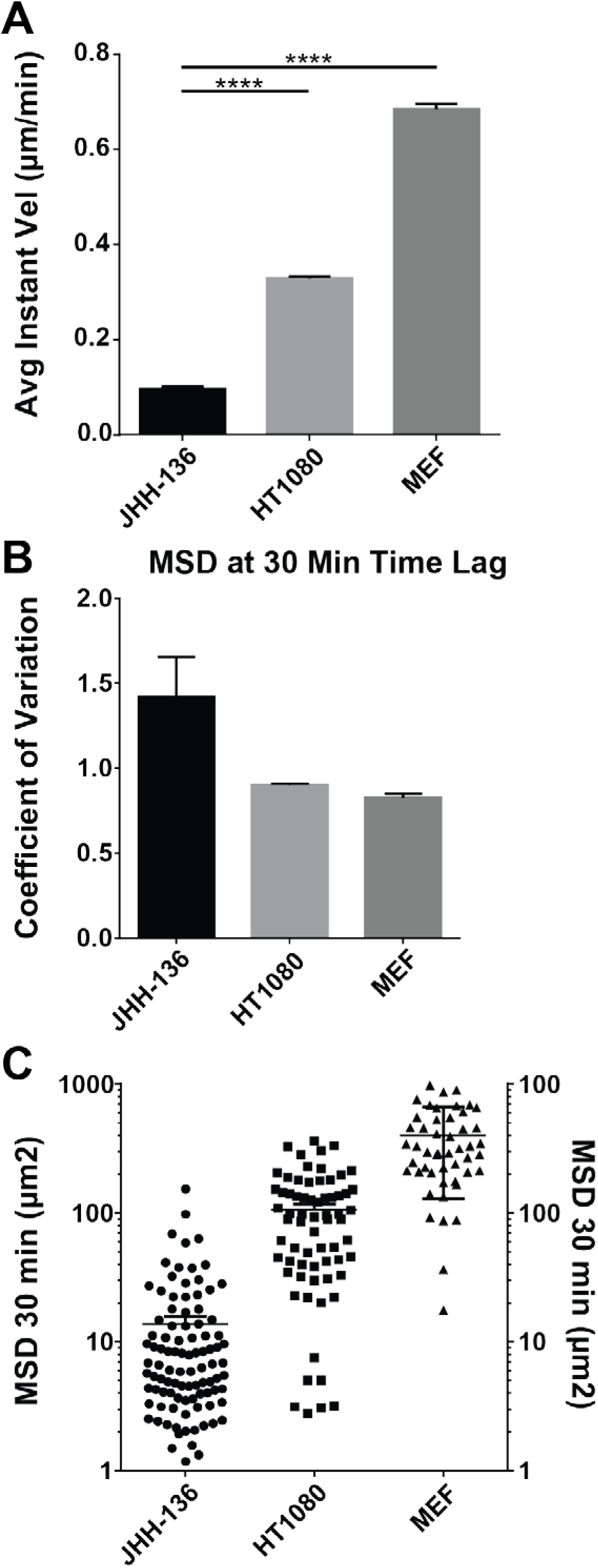
Comparison of suspension GBM oncosphere cells in brain tissue slices to HT1080 cells in 3D and MEF cells in 2D. (A) Population-averaged instantaneous velocities at 10 min time lags comparing JHH-136 suspension GBM oncosphere cells to two other conditions: HT1080 cells in 3D collagen gels and MEF cells on a 2D collagen coating. (B and C) Comparison of coefficient of variation (B) and dot plots (C) of MSD values at 30-min time lag. JHH-136 cells demonstrated a greater spread of values, highlighting the strong degree of heterogeneity found in the GBM tumor cells.

**Supplementary Figure 2.**
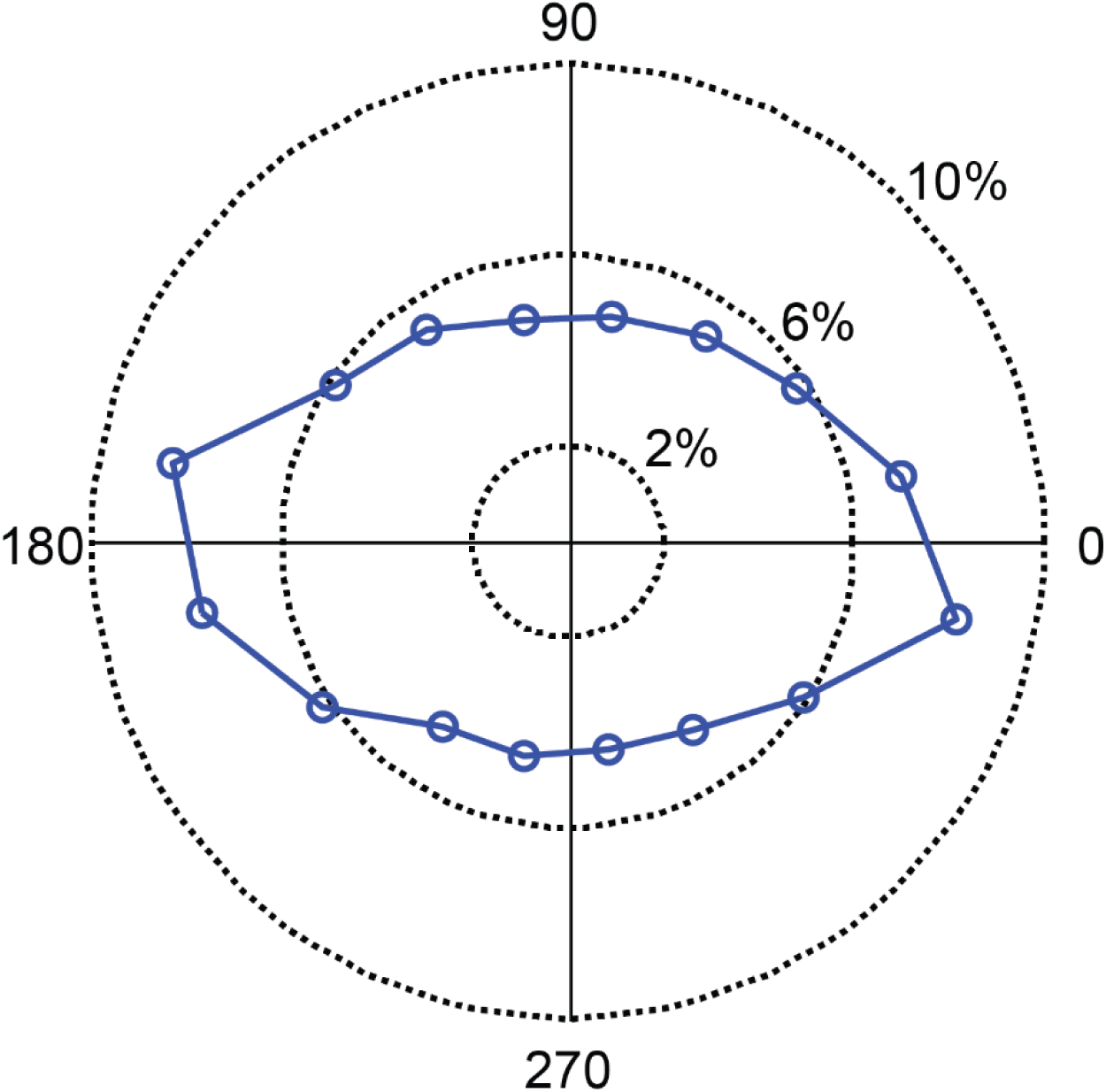
Trajectory frequency distribution showing an anisotropic pattern of movement.

**Supplementary Figure 3.**
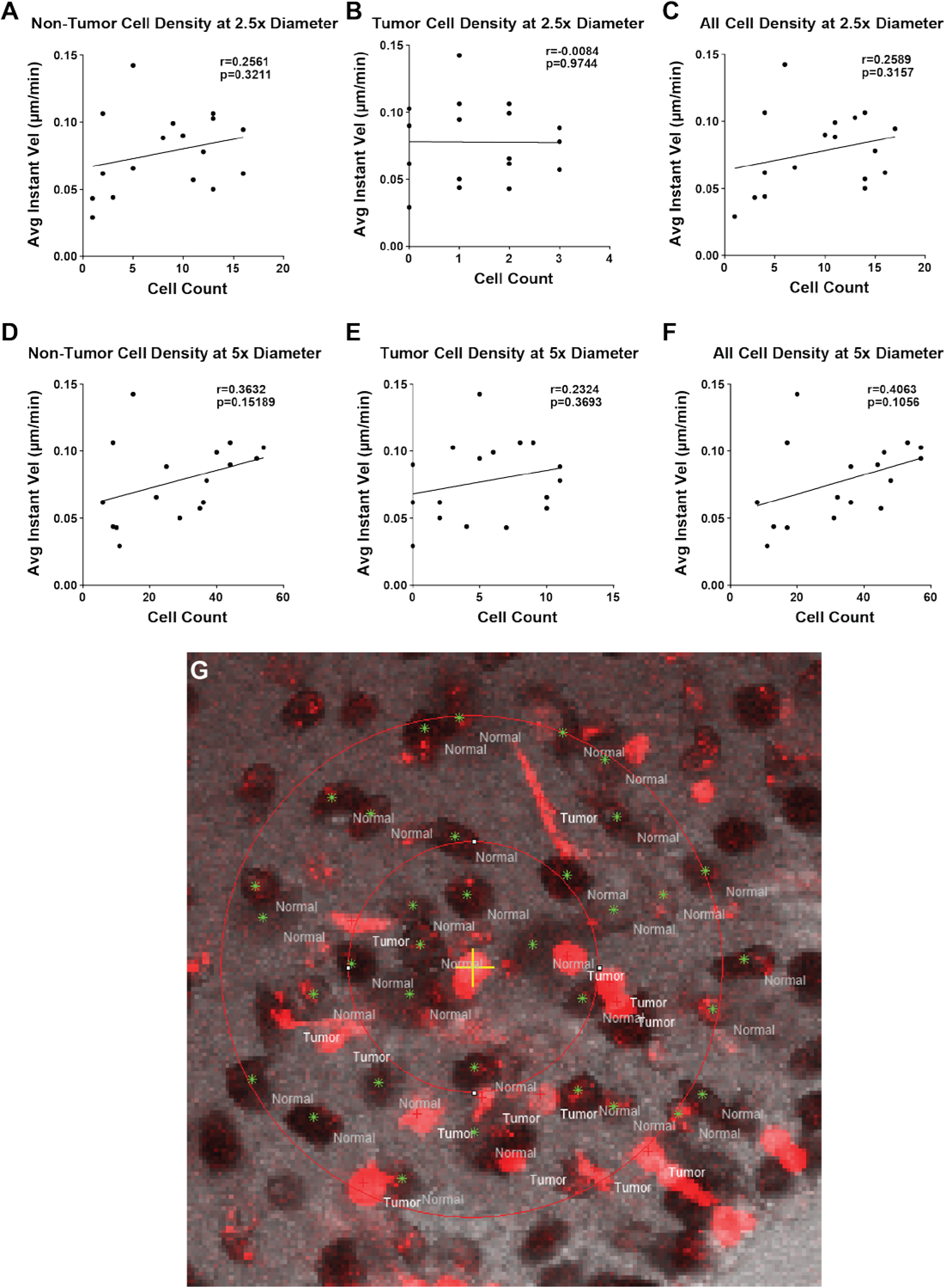
Tumor cell density and normal cell density do not display a significant correlation with population-averaged instantaneous velocity. (A-C) Non-tumor (A), tumor (B), and total (C) cell densities in a 40.85 µm diameter circle. Non-tumor cell, tumor cell, and overall cell densities demonstrated no correlation with population-averaged instantaneous velocities. (D-F) Non-tumor (D), tumor (E), and total (F) cell densities in an 81.7 µm diameter circle, showing the same correlative relationships as the smaller diameter densities. (G) Differential interference (DIC) and fluorescence confocal imaging showing physiologic parenchyma (DIC) and tRFP-labeled tumor GBM tumor cells demonstrating cell densities. Scale bar, 50 µm.

**Supplementary Figure 4.**
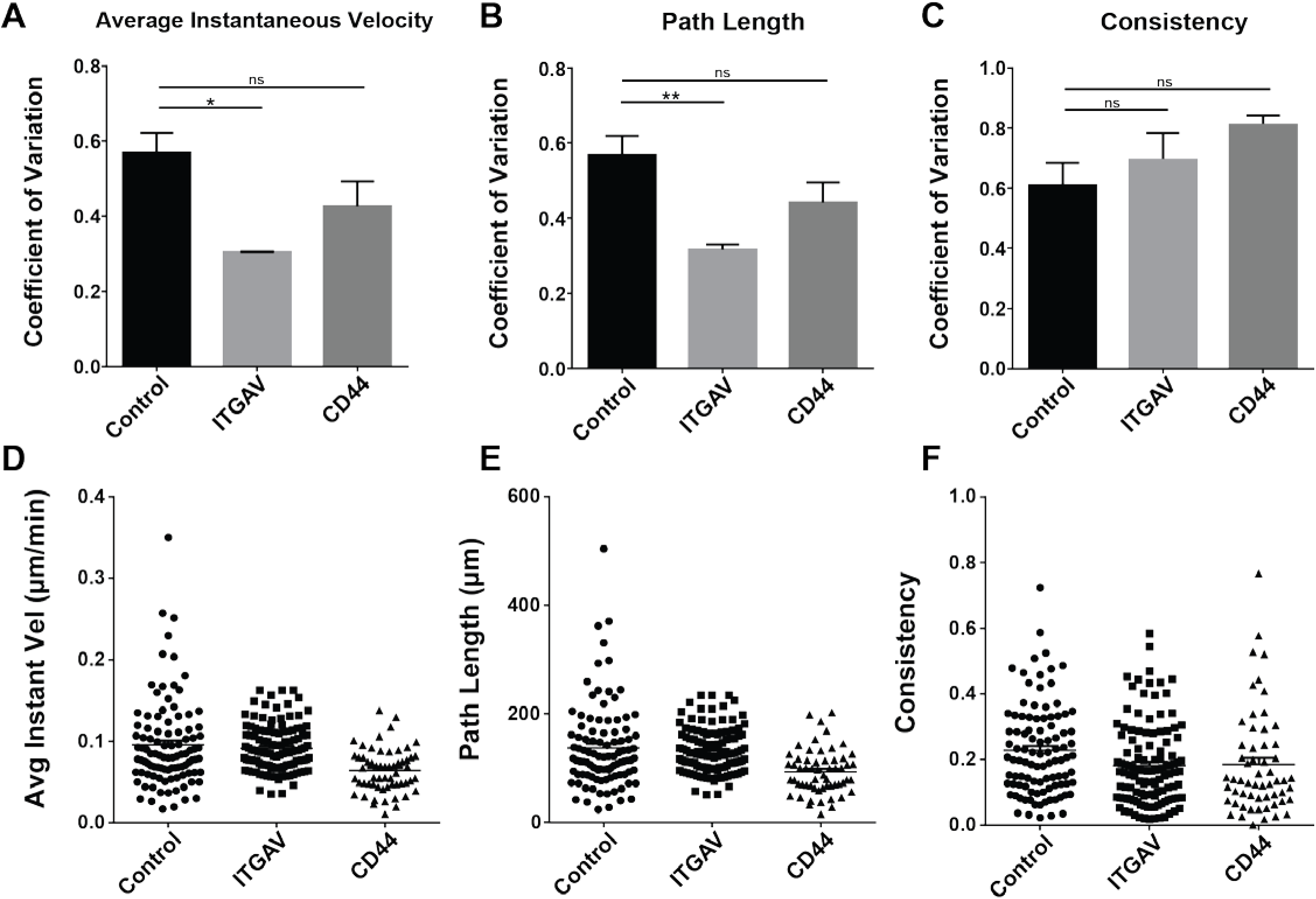
Cell population heterogeneity is not significantly reduced with depletion of either ITGAV or CD44. (A-C) Coefficient of variation for Control, ITGAV-depleted, and CD44-depleted cells in (A) population-averaged instantaneous velocity, (B) total path length, and (C) consistency as measured by final distance from origin divided by total path length. (D-F) Dot plots showing the population heterogeneity found in the cell motility characteristics. (D) Population-averaged instantaneous velocity, (E) total path length, and (F) consistency. ns: non-significant; *: p ≤ 0.05; **: p ≤ 0.01.

